# Fragile dynamics enable diverse genomic determinants to influence arrhythmia propensity

**DOI:** 10.1101/101162

**Authors:** Tim J Kamerzell, Eric A Sobie, Kai-Chen Yang, Jeanne M Nerbonne, Calum A MacRae, Ravi Iyengar

**Author notes:** Current address: Department of Pharmacology, National Taiwan University School of Medicine, Taipei, Taiwan (Kai-Chen Yang). Correspondence to: Ravi Iyengar. Department of Pharmacology and Systems Therapeutics and Systems Biology Center New York, Icahn School of Medicine at Mount Sinai, New York City, NY 10029.

## Abstract

Genotype-phenotype relationships are determinants of human diseases. Often, we know little about why so many genes are involved in complex common diseases. We hypothesized that this multigene effect arises from the relationship between genes and physiological dynamics. We tested this hypothesis for arrhythmias as physiological dynamics define this disease. We integrated graph theory analysis of genomic and protein-protein interaction networks with dynamical models of ion channel function to identify the physiological dynamics of genome wide variation for five different arrhythmias. Regulatory networks for the cardiac conduction system and arrhythmias were constructed from GWAS and known disease genes. Electrophysiological models of myocyte action potentials were used to conduct extensive parameter variations to identify robust and fragile kinetic parameters that were then, using regulatory networks, associated with genomic determinants. We find that genome-wide determinants of arrhythmias that represent many cellular processes are selectively associated with fragile physiological dynamics of ion channel kinetics. This association predicts disease propensity. Deep RNA sequencing from human left ventricular tissue of arrhythmia and control subjects confirmed the predictive relationship. Taken together these studies show that the varied multigene effects of arrhythmias arises because of associations with fragile kinetic parameters of cardiac electrophysiology.

**Significance Statement:** Our understanding of the genetics of common diseases has advanced exponentially over the past decade. We now know that differences and variation in multiple genes contribute to disease susceptibility with significant heterogeneity in the phenotype. However, how genetic variation contributes to disease phenotypes remains unknown. We hypothesized that the relationships between physiological dynamics and genetic architecture is a fundamental determinant of disease susceptibility and genetic heterogeneity. To test our hypothesis, we integrated mathematical models of cardiac electrophysiology with genetic network models of cardiac arrhythmias. We found that disease related genome variants were selectively associated with fragile kinetic parameters that predict disease propensity and identified several novel cellular processes associated with arrhythmogenesis.

## INTRODUCTION

The genetic architecture of many common diseases has been extensively studied over the past decade providing valuable insight into the etiology of disease (1, 2). Differences and variation in genetic architecture contribute to disease susceptibility (1–4). Despite tremendous advances in the genetic basis of these diseases our understanding of how and why genomic changes manifest as disease phenotype remains incomplete and our ability to predict disease risk is often inadequate. This is especially true for many complex progressive diseases including autoimmune disorders, neuropsychiatric diseases, diabetes and cardiovascular diseases. The relationships between genotype and phenotype are complex because they are based on genetic heterogeneity, the environment and the dynamics of physiology resulting in nonlinear interactions between genes and gene products (1, 2, 5–7). We sought to identify a simple principle as to why so many varied genomic determinants are involved in disease phenotype. We hypothesized that the involvement could be due to a selective association with fragile features of the physiological dynamics. To test this hypothesis we used a model integration approach combining network models with multi-compartment ODE models.

Arrhythmias are complex diseases that most often arise from mutations in genes that encode ion channels ultimately leading to altered physiological dynamics. Nevertheless, recent evidence suggests that drivers of conduction abnormalities and arrhythmias can extend well beyond mutated channel proteins. For example, Wnt11 has been associated with the ECG PR interval in GWAS (8) and in a separate study was shown to pattern a myocardial electrical gradient through regulation of the L-type Ca^2+^ channel (9). Similarly, the transcription factor PITX2, has been associated with atrial fibrillation in GWAS (10) and was subsequently shown to regulate ion transport and impair calcium handling (11, 12). A multi-model approach that integrates physiological dynamics, cell regulatory networks and genetic architecture could provide insight as to why such varied genes are involved. Using canonical cell biological knowledge it is possible to schematically describe the relationships between cell regulatory networks to cardiac myocyte action potentials and consequently to conduction abnormalities. To mechanistically connect genomic variations to physiological dynamics we need to integrate two types of computational models: (1) network models that delineate the topology of regulation in terms of gene products (proteins) and their interactions and (2) dynamical models that describe the time course of physiological responses (i.e. ion channel function) in terms of protein function. Both types of models have been used to study arrhythmias. Previous network analysis studies have shown considerable overlap in the human interactome between products of genes that can cause Long QT syndrome when mutated and targets of drugs that can cause adverse cardiac events (13, 14). This analysis generated the prediction, subsequently validated by clinical observations, that loperamide could cause arrhythmias (15). A different computational approach, combining statistical regression analysis with dynamical models of cardiac electrophysiology, at the cellular and organ levels, also provided deep insight into the quantitative dynamic relationship between the cardiomyocyte action potential and the heart’s electrical activity (16). Systematic parameter sensitivity analysis of cardiomyocyte models generate precise predictions of how alterations in any cardiac ionic current may either exacerbate or mitigate pathological alterations to the electrocardiogram (16–20). This approach can quantitatively classify electrophysiological parameters as either fragile, meaning that small changes in parameters can cause large physiological alterations, or robust, meaning that cells or tissues can tolerate large changes to the parameter values without altering physiological responses.

To test our hypothesis that disease related genes might associate with the fragile features of physiological dynamics we integrated the network and dynamical models. We found that disease related genome variants were selectively associated with fragile kinetic parameters. Model integration also identified several cellular processes that could be associated with arrhythmogenesis. Using deep RNA sequencing data from human cardiac tissue from patients with arrhythmias we verified many key predictions from our integrated model including an association of arrhythmias with fragile kinetic parameters and the involvement of pathways such as propanoate metabolism. We combined multiscale data with unsupervised learning to identify genomic determinants of disease that correlate with fragile genes allowing us to demonstrate why genes belonging to many cellular processes are associated with increased disease propensity.

## RESULTS

We studied the systems biology of arrhythmias through integration of graph theory and dynamical modeling, combined with transcriptomic analysis of human heart tissue, to better understand how genomic architecture can contribute to cardiac electrical phenotypes. Figure 1 shows a schematic of the computational and experimental methods used in this study. Initially, we compiled prior information from classical disease gene discovery and GWA studies to comprehensively catalog genomic determinants, including mutations, SNPs and other genomic variants, known to be associated with ECG characteristics and five types of arrhythmias. This database of genes was then used to build protein-protein interaction networks and identify potential novel disease mechanisms. Next, we used well known ordinary differential equation based models of the cardiomyocyte action potential to identify parameters (kinetic properties of gene products) that were either sensitive (fragile) or robust to modulation. Our approach surprisingly identified that fragile kinetic parameters are preferentially associated with disease related genomic determinants. The network analysis also generated the unexpected prediction that arrhythmia genes are closely associated with genes involved in previously unrelated cellular processes, including propanoate metabolism, the ubiquitin proteasomal degradation pathway, and histone modification and chromatin remodeling. We then validated numerous predictions from our models using transcriptomic profiling of human left ventricular tissue from patients with arrhythmias.

**Figure 1.**
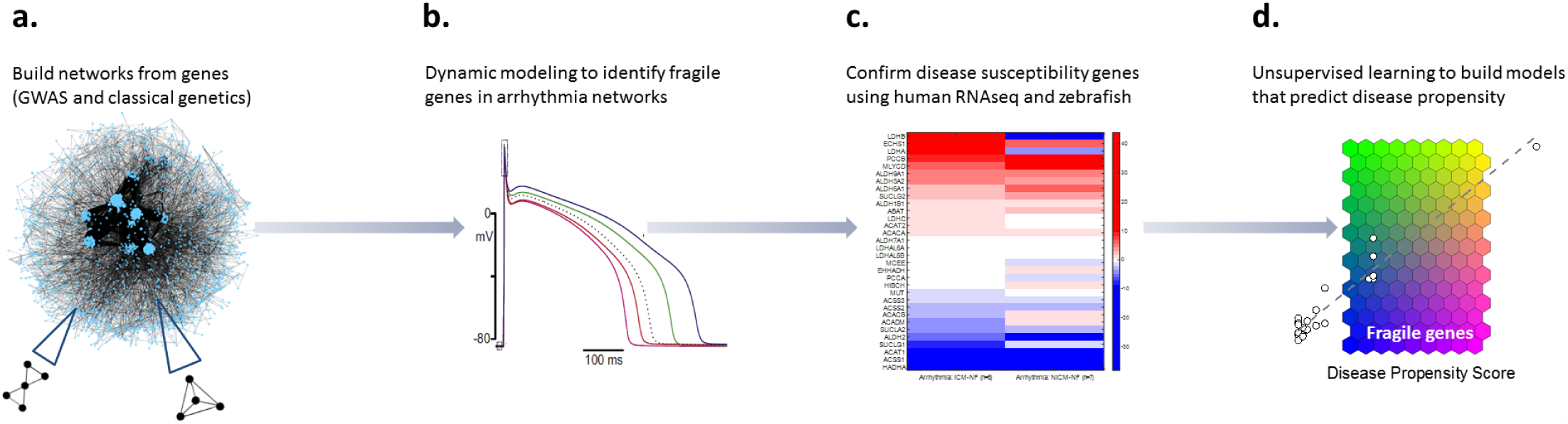
Schematic of methods used in this study. (a) Networks of ECG components and arrhythmias were constructed from all known genes identified in genome wide association studies (GWAS) and classical genetic approaches. Biological interaction networks identified potential novel disease mechanisms. (b) Ordinary differential equation models of the cardiac action potential were used to identify genes with sensitive changes to model parameters and subsequently mapped to genes within the arrhythmia networks. (c) Human RNA sequencing verified several key predictions from the arrhythmia network and ODE dynamic simulations. (d) Various unsupervised learning approaches were used to identify genomic association with disease genes that predict disease propensity.

### Networks from ECG and arrhythmia genomic determinants identify new biological associations

We utilized published human studies to identify all known genes associated both with the ECG and five different arrhythmias: atrial fibrillation, QT-Opathies, Brugada, catecholaminergic polymorphic ventricular tachycardia (CPVT) and progressive cardiac conduction defect (PCCD) **(Supplementary Fig. 1a-b)**. We identified relevant genes from classical disease gene discovery studies and mapped or reported genes from SNPs identified in genome wide association studies (GWAS). The full list of seed genes (153 for ECG and 109 for arrhythmia networks) is provided in **Supplementary Tables 1-2**. Often the same gene/gene-products are involved in multiples phases of the ECG or different types of arrhythmias. These relationships are shown in **Supplementary Figures 1c-e**. Individual genes may contribute to multiple electrophysiological responses (pleiotropy) and multiple genes can affect individual phenotypes (epistasis) (21, 22). Examples include, SCN5A, the gene encoding for the cardiac voltage-gated sodium channel, associated with several ECG characteristics (PR interval, QRS duration and QT interval) and arrhythmias (atrial fibrillation, QT-opathies, Brugada syndrome and PCCD) (8, 21, 23–25), and cardiac transcription factors such as NKX2-5, SOX5, and TBX5. Many of the genes are known ion channels or channel associated membrane proteins, but other known functions include cardiovascular development and intracellular signaling. The overlap of individual ECG and arrhythmia network genes is shown as Venn diagrams in **Supplementary Figure 2**.

Using genes associated with particular arrhythmias and ECG characteristics as seed nodes, we developed biological interaction networks by expanding around these nodes using protein-protein interactions (26, 27). Networks were constructed using direct protein interactions with seed genes and one intermediate node to connect the seed nodes. In the Supplementary Materials Section we describe the databases from which human protein interactions were imported and the methods used to expand ECG and arrhythmia network genes to 4190 and 4955 components, respectively. Representative networks for five separate arrhythmias and components of the ECG are shown in **Supplementary Figure 3**.

Atrial fibrillation is the most common human cardiac arrhythmia with a rapidly increasing prevalence and incidence worldwide. The atrial fibrillation network is shown in Figure 2a with several highlighted clusters of densely connected nodes identified based on connectedness and biological function. For example, the cluster of red nodes (genes) in the top right corner of Figure 2 is enriched for genes involved in propanoate metabolism. Functional gene enrichment of each arrhythmia network was used to identify biological processes potentially contributing to disease. Overrepresented gene ontology terms were determined from lists of genes that make up each network (28). Figure 2b shows a functional enrichment network of the most significant biological processes involved with atrial fibrillation. Nodes in the enrichment network indicate the most over-represented biological processes, with node size scaled using the betweeness centrality and color coded according to the biological process (metabolism, histone modification, etc.). Edges between nodes, with thickness scaled using edge betweeness, indicate that multiple genes contribute to each process. For example, genes within the atrial fibrillation network that are involved with Wnt signaling also contribute to heart development, chromatin modification and androgen signaling, among others. **Supplementary Table 3** summarizes the most significant enrichment terms used to describe the set of network genes for all five arrhythmias.

**Figure 2.**
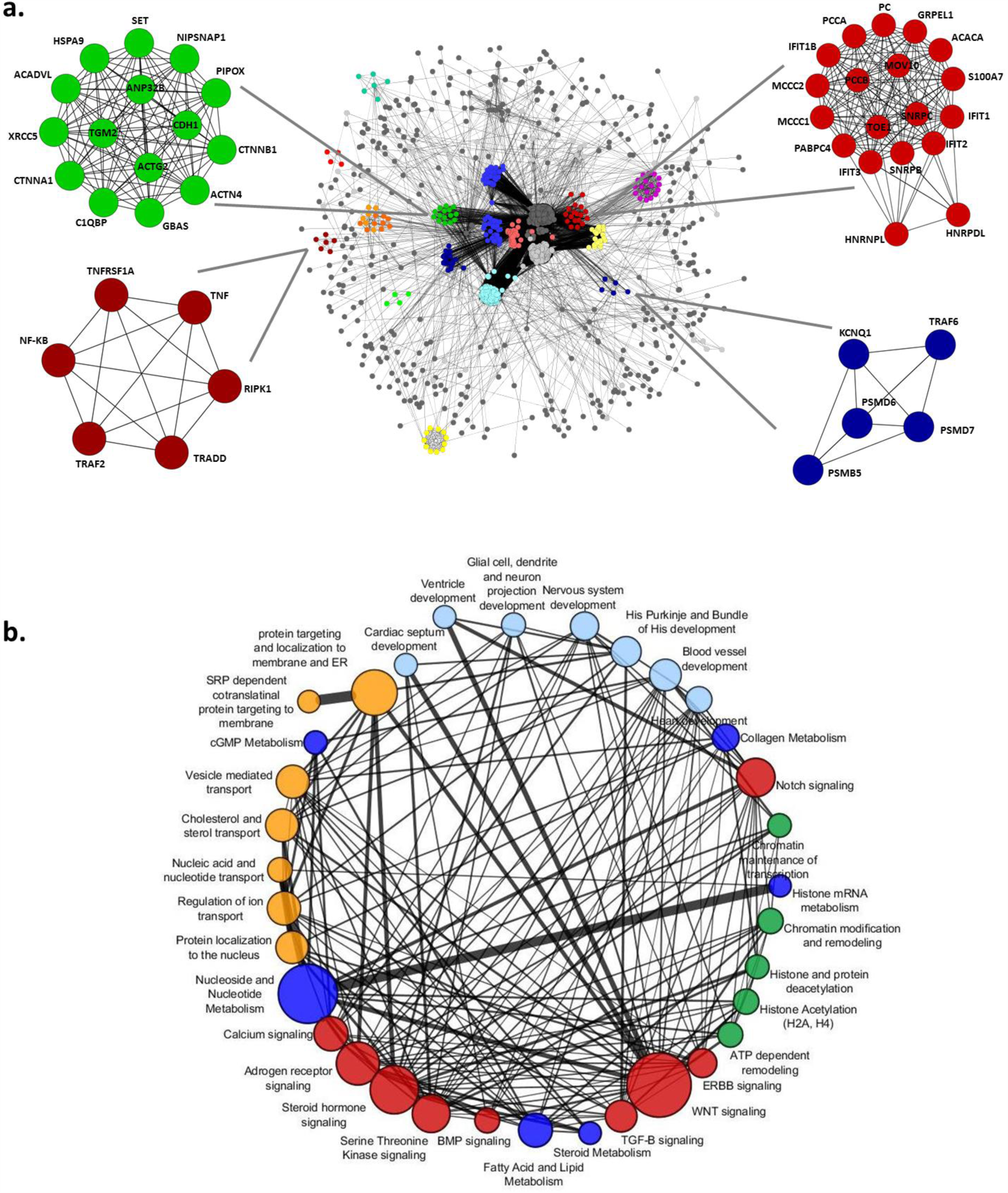
The atrial fibrillation network. **(a)** Representative atrial fibrillation interaction network. The nodes in each network represent a single protein or gene and the edges connecting two nodes indicate an interaction. Clusters of highly interconnected nodes and regions in the network are shown in color. Several clusters are magnified to highlight specific connections and the associated genes within that cluster are labeled. Clusters of genes associated with propanoate metabolism (red node cluster in top right corner), protein ubiquitination (blue node cluster), apoptosis and protein processing (red node cluster in lower left corner) and cell-cell junctions and protein localization to the surface (green node cluster) are shown. **(b)** Functional gene enrichment network highlighting the most significant biological processes identified from the atrial fibrillation arrhythmia network genes. Node size is scaled using the betweeness centrality and color coded according to the associated biological process. Edges between nodes are scaled using edge betweeness and indicate a significant number of genes contributing to the biological processes of both connected nodes. For example, genes within the AF network involved with Wnt signaling are also involved with heart development, chromatin modification, androgen receptor signaling, etc. See **supplementary table 3** for a summary of various molecular processes associated with functional gene enrichment of each arrhythmia network. The table is color coded by biological process to match the nodes and network in figure 2D1.

The biological functions of genes contributing to the arrhythmia networks can be classified into seven major groups; cellular signaling, transport (including ion channels), metabolism, protein turnover, gene expression, chromatin and histone modifications, and development. The similarities and differences between arrhythmia networks were identified and categorized for each of the seven major groupings (**Supplementary Table 3**). Significant overlap of enriched biological processes from genes involved with arrhythmias was found. This analysis revealed several cellular processes that have not been previously associated with arrhythmogenesis. For example, RNA splicing, ubiquitin degradation, membrane transport and metabolism, more specifically propanoate metabolism, were all enriched in multiple arrhythmia networks. The network models therefore generate potentially testable hypotheses by suggesting that these biological processes may be related to the pathophysiology of arrhythmias. Although the expanded networks most certainly contain some genes that do not contribute to disease it is less likely that these “non-disease” genes will be identified as highly enriched or components of highly connected dense clusters. Furthermore, we verified many of the biological processes identified in the networks using human RNA sequencing and functional knockdown experiments in zebrafish.

The macro-scale topology of arrhythmia networks shows well-connected communities linked to more isolated communities. Each network is defined by a number of dense communities or hubs connected to other regions of the network via less well connected or sparse clusters. The Venn diagrams readily highlight the substantial genetic overlap and number of unique genes between the various ECG and arrhythmia components suggesting a large number of potential disease or modifier genes involved with the cardiac electrical response including the diversity of molecular responses leading to phenotypic heterogeneity. That is, there are many different routes to disease. To further characterize network structure we calculated topological parameters and various network statistics (**Supplementary Figures 4-13**). Both the ECG and arrhythmia networks are approximately scale free with power law like degree distributions (**Supplementary Figures 4 and 6**). The degree distributions of these real world genetic interaction ECG and arrhythmia networks systematically and expectedly deviate from the degree distributions of random networks with the same number of nodes and edges (**Supplementary Figures 5 and 7**). To assess global (average) network metrics, we calculated the average clustering coefficient, density, heterogeneity, centralization and characteristic path length of each network (**Supplementary Figures 8 and 9**) (29). The network statistics confirmed the visual impression that these networks are highly connected, dense and heterogeneous with high degrees of centrality measures as compared to random networks. These findings are expected and suggest that arrhythmia networks comprise a large number of essential disease genes or modifier disease genes with potentially fragile interdependencies (30–32).

### Deep RNA sequencing of human subjects with arrhythmias confirms association of new cellular processes with disease

We used deep RNA sequencing of human subjects with arrhythmias to independently confirm the novel cellular processes associated with arrhythmias that were separately identified from enrichment analysis of the biological interaction networks. The ventricular myocardial transcriptional profiles of human subjects with arrhythmias were analyzed and compared with those of human subjects without disease (Figure 3). Genes with the largest difference in expression were statistically analyzed for significance and those with statistically significant differences (p ≤ 0.05) were included in gene enrichment analysis. The Fisher exact test with associated p-values was used for enrichment analysis in the program Enrichr (28). Many of the same biological processes identified from enrichment analysis of the biological network models were also identified using enrichment analysis of the genes from human RNA sequencing. Representative heat maps of differences in expression profiles involved with propanoate metabolism (Fig. 3a) and membrane transport, RNA splicing and cardiovascular development (Fig. 3b) highlight some of the changes in the respective cellular processes enriched in human subjects with arrhythmias. These findings confirm the identification of novel cellular processes associated with arrhythmias. In addition to identifying new cellular processes, previously known mechanisms of arrhythmogenesis were also confirmed. A comparison of overlapping gene enrichment terms identified using network analysis and human RNA sequencing is shown in Figures 3c-d highlighting many new and well known biological processes. In fact, the overlap of all human RNA seq enrichment terms with the atrial fibrillation network enrichment terms is nearly 70% and greater than 50% overlap is observed with all other arrhythmia networks (Fig. 3d).

**Figure 3.**
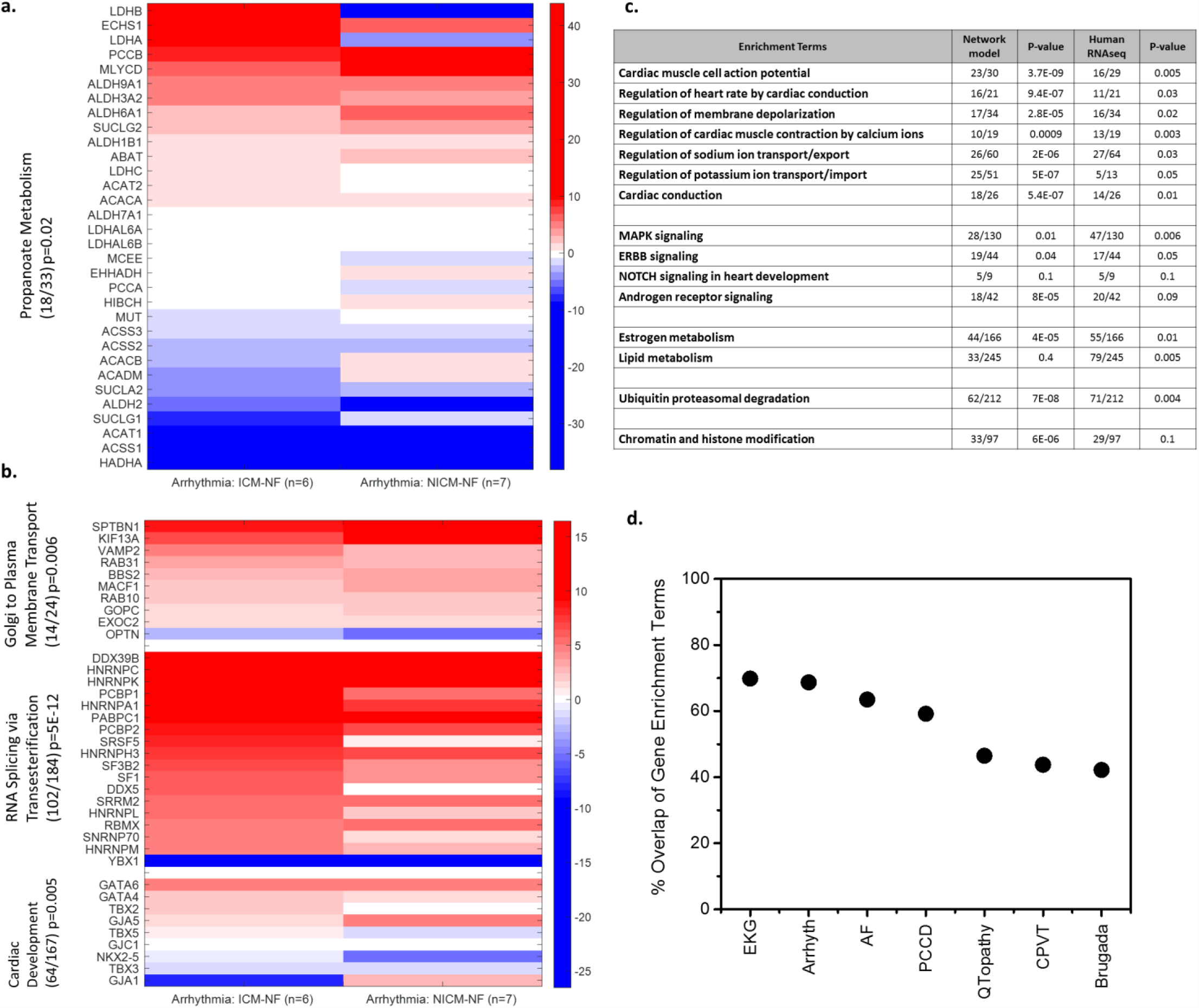
Human RNA sequencing reveals expression level differences and confirms biological processes associated with arrhythmias. Representative heat maps of gene expression level differences between human subjects with arrhythmias and without arrhythmias. **(a)** Heat map of expression level differences in key propanoate metabolism genes. **(b)** Heat map of expression level differences in key biological processes determined to be important by gene enrichment analysis. **(c)** Table of highly ranked biological process terms using gene enrichment and gene ontology. The table compares the top biological processes determined using network models and human RNA sequencing data. The p-value is calculated using the Fisher exact test which is a proportion test that assumes a binomial distribution and independence for probability of any gene belonging to any set. Gene enrichment analysis and statistics were calculated using the Enrichr program previously described by Chen *et al* (28). **(d)** Overlap of gene enrichment terms comparing network models to human RNA sequencing. The statistically significant gene enrichment terms were identified from the genes in human RNA sequencing and separately for each individual arrhythmia or ECG network. The number of terms that overlap were calculated as a percentage. **ABBREVIATIONS**: ICM-ischemic cardiomyopathy; NICM-non ischemic cardiomyopathy; NF-control subjects without arrhythmias or cardiomyopathy.

### From topology to dynamics: Mapping the kinetic parameters that contribute to the fragility of dynamics onto genes

The topological network and enrichment analyses are useful in understanding at a global level the subcellular processes involved in arrhythmias, however, network models do not provide information about physiological dynamics. Hence we sought to integrate dynamical models of cardiac electrophysiology with the gene interaction network models. This integrated computational analysis can help us ascertain the value of genomic changes in altering physiological responses. The dynamical models identify both fragile and robust genes potentially contributing to disease which may also be useful for predicting disease propensity.

Using well-established dynamical models, we conducted massive simulations of atrial, ventricular and SA node cardiac action potentials by systematically varying the parameters that define the conductances and rates of ion transport (16, 18, 33–36). Figures 4a-c show representative voltage, intracellular calcium and subspace calcium tracings from a single simulation of the ventricular action potential. Figures 4d-f shows many ventricular, atrial and SA node action potential tracings from massive simulations while varying the model parameters that define ionic conductances. Changing kinetic parameters differentially modulates the voltage tracings. Some kinetic parameters substantially influence the voltage tracings while others only modestly perturb voltage. These simulations allowed us to quantify how sensitive the different action potential models are to changes in the various ionic conductances which can then be linked to genes.

**Figure 4.**
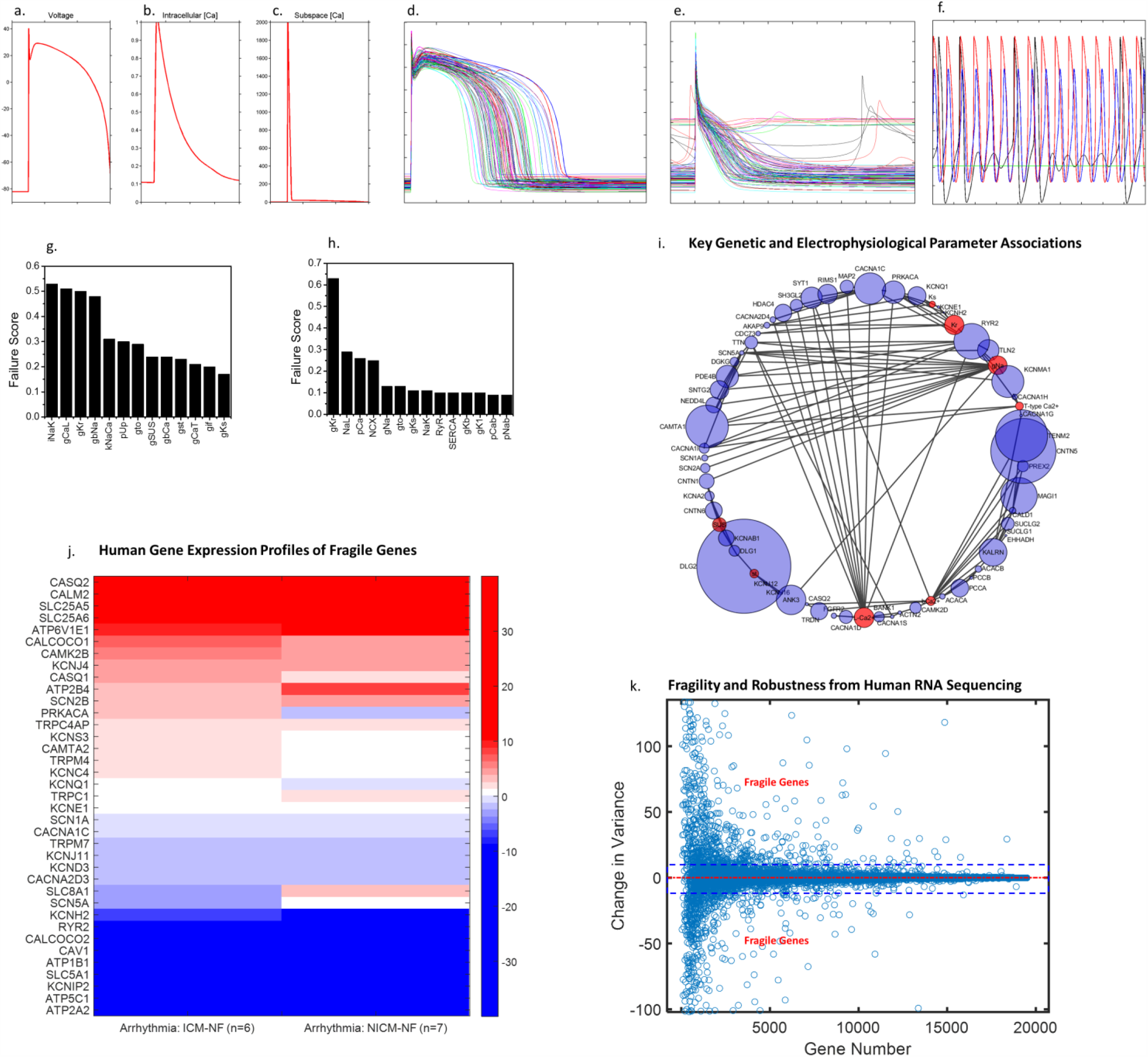
Parameter sensitivity and failure analysis reveals fragile electrophysiological parameters that underlie arrhythmogenesis are also associated with disease genes from human RNA sequencing. Representative **(a)** voltage, **(b)** intracellular calcium and **(c)** subspace calcium tracings from a single simulation of the ventricular cardiac action potential. Many representative **(d)** ventricular, **(e)** atrial and **(f)** SA node action potential tracings from large scale simulations while varying the models parameters. Dynamical model parameter failure scores, shown as bar graphs, calculated from large scale simulations of the **(g)** ventricular and **(h)** atrial action potential. Failure scores represent the sensitivity of the cardiac electrophysiological response to individual parameter variation. Larger values of the failure score indicate a greater sensitivity to perturbation and an increased propensity for arrhythmogenesis. For bar graphs of regression coefficients from parameter sensitivity analysis of large scale simulations see **supplementary figures 14** and **15**. **(i)** Fragile parameter-gene network showing the connection of key fragile genes to model parameters. Red nodes indicate model parameters and blue nodes indicate linked genes. Node sizes are scaled to represent electrophysiological parameter fragility and genetic variation. A list of model parameters and several associated key fragile genes included in the gene networks listed with their corresponding failure scores is shown in **supplementary figure 16**. **(j)** Representative heat map of several fragile gene expression profiles of human subjects with arrhythmias. The change in gene expression between human subjects with arrhythmias compared to control (subjects without arrhythmias) are shown as red (positive) or blue (negative). **(k)** Plot of the change in variance between human subjects with arrhythmias and control subjects for 20,000 different genes using human RNA sequencing data. The mean change in variance is shown as a dashed red line. The 90^th^ percentile of values are shown as dashed blue lines. All fragile genes lie above or below the dashed blue lines.

Multivariate regression and failure score analysis were used to quantify the sensitivity of the action potential duration and calcium transients to changes in individual and pairwise parameters (Figures 4g-h **and Supplementary Figures 14-15**). To perform pairwise parameter analysis we simultaneously varied two parameters while recording the outputs at the end of the simulation. The magnitude of the regression coefficients from regression analysis are quantitative measures of the sensitivity of simulation outputs (e.g., voltage, calcium transients) to parameter perturbation (e.g., varying L-type Ca^2+^ conductance). We also calculated a catastrophic failure score that represents the relative likelihood of a failed action potential due to changes in kinetic parameters (ionic conductances). Figures 4g-h show the calculated failure scores for ventricular and SA node models while varying each parameter. The failure scores and regression coefficients (parameter sensitivities) represent quantitative dynamic measures of the cardiac electrophysiological response to perturbation. We defined fragile parameters as those that contribute to nearly 80% of the failures of action potential propagation upon perturbation. Thus the molecular kinetic parameters most sensitive to perturbation are more fragile at a whole cell and organ level as these parameters are more likely to contribute to action potential failure and arrhythmogenesis when perturbed even by a small amount.

Our analysis shows that the cardiac action potential is most sensitive to perturbations in the L-type Ca^2+^ current (G_CaL_), Na^+^-Ca^2+^ exchange current (kNaCa, NCX), and rapid delayed rectifier K^+^ current (G_Kr_). The Na^+^ current, Na^+^-K^+^ pump current (I_NaK_) and the cardiac ryanodine receptor channels (RyR) current are also sensitive parameters. The relative importance of each parameter varies according to the model used (SA node, ventricular or atrial AP); however, nearly 80% of the failure scores can be accounted for by sensitivity to perturbation in the aforementioned parameters. Multiparameter sensitivity analysis thus provides a quantitative measure of simultaneous electrophysiological perturbation that can be connected phenomenologically to genes comprising arrhythmia networks and used to help explain phenotypic variation.

To determine the relationship between genomic characteristics to electrophysiological dynamics and phenotypes we mapped genes identified in the networks to the sensitive model parameters. Network genes were associated with the dynamical model parameters based on function of the gene and parameter. **Supplementary Figure 16** lists some of the important associations between dynamical parameters and network genes. We refer to the nodes that map to the most sensitive parameters as fragile genes – those that, when altered, might result in a greater propensity for developing arrhythmias. An additional network was then constructed connecting dynamical model parameters to genes with the node size scaled to represent parameter fragility and genetic variation obtained from the NCBI databases and Variation Viewer (Fig. 4i). For example, the Na^+^-Ca^2+^ exchange current, which directly maps to the network node SLC8A1, was shown to be a sensitive parameter from the action potential simulations. Similarly, KCNQ1 and KCNE1, known potassium voltage-gated channel genes required for repolarization, directly map to the slow delayed rectifier K^+^ current parameter, G_Ks_.

### Human RNA sequencing supports the association of genes with fragile dynamics with arrhythmias

Having computationally identified that specific genes involved with arrhythmias have fragile dynamics we sought to test if this computational prediction could be experimentally validated. Fragile dynamics means sensitivity to perturbation and can arise both from variation in kinetic parameters and levels of the reactants. Assuming that mRNA levels could overall reflect protein levels we formulated the hypothesis that fragile genes would show greater changes in disease state or would result in greater changes in other protein levels as a compensatory change. This is in fact how we define sensitive parameters in our dynamical model. Therefore, using human RNA sequencing as our experimental approach and genomic analysis, we sought to determine if this association was valid and identify new mechanisms potentially contributing to disease. In these analyses we also sought to determine the characteristics and potential mechanisms underlying this association.

We evaluated human ventricular tissue gene expression profiles of the genes computationally identified to have fragile dynamics in subjects with and without arrhythmias (Figure 4j). The methods used for RNA sequencing have been previously described (21) and in the Supplementary Materials section. Figure 4j shows the difference in expression of select fragile genes (as a heat map) between subjects with and without arrhythmias, highlighting the substantial changes when a perturbation (human subjects with arrhythmias) is introduced. The Na^+^-K^+^ ATPase gene has the highest failure score based on the computational ventricular model and also shows one of the highest changes in gene expression in human ventricular tissue in arrhythmia patients. There are considerable differences in expression levels of many genes with fragile dynamics. This data supports our hypothesis that genes with fragile dynamics are likely to show greater changes in their tissue level expression in the disease state.

We also used the human RNA expression profiles to further support the association of fragile dynamics with arrhythmia genes by calculating *in-vivo* robustness and fragility of genes using previously described methods based on variational principles (37). When using human RNA sequencing, fragility can be defined as large changes in gene expression variation in the presence of perturbation (arrhythmogenesis) and/or the change in variance of gene expression between subjects with and without arrhythmias. The variance of robust genes is predicted not to change a significant amount between control subjects and those with disease. If both control and disease gene expression levels are highly variable then the change in variance will be small. Similarly, if the variance of both control and disease gene expression levels is small then the change in variance will be small. In contrast, if the variance of control genes is small but the variance of disease genes is large (vice versa) then the change in variance will be large. That is, fragile genes may show large changes in variance upon perturbation suggesting sensitivity to changes in disease state.

Figure 4k shows the change in gene expression variance between human subjects with arrhythmias and subjects without arrhythmias for nearly 20,000 different genes. The 90^th^ percentile of change in variance values are between the two dashed blue lines outlined in the figure. That is most genes are characterized by relatively small changes in variance. These genes with low change in variance can be considered robust genes as previously described (37). The red dashed line denotes the median value of change in variance for all 20,000 genes. The change in variance, of all genes with previously determined fragile dynamics by our computational methods, is greater than the 90^th^ percentile and most values are greater than the 95^th^ percentile suggesting these genes whose proteins have fragile dynamics are highly sensitive to changes in the disease state. The change in variance of three propanoate metabolism genes (HADHB, LDHA and LDHB) and four genes involved in calcium regulation and current (KCNIP2, ATP2A2, PLN and CASQ2) have values greater than the 99^th^ percentile. Therefore, human RNA sequencing of subjects with and without arrhythmias independently provides strong support for the computational finding associating fragile genes with disease.

### Unsupervised learning identifies correlation between genomic architecture and fragile genes in subjects with arrhythmias

We next tested whether different types of genomic variations selectively contribute to the fragile dynamics of disease genes and provide additional mechanistic insight contributing to the likelihood of developing disease. To identify the genomic characteristics contributing to the fragility of the products of arrhythmia genes, we utilized two unsupervised learning methods, Self-Organizing Maps (SOMs) and Principal Component Analysis (PCA), to identify and classify the genomic differences (SNPs, CNVs, CpG counts, etc.) between clusters of fragile and robust genes comprising arrhythmia networks (Figure 5 **and supplementary Figure 17-18**). SOMs are dynamical, adaptive, nondeterministic and nonlinear, and can help to identify emergent properties. We trained the SOMs using diverse genomic characteristics (SNPs, CNVs, CpG counts, etc.) of the arrhythmia networks (**Supplementary Table 4**). Figure 5a shows a heat map of genomic characteristics for individual genes used to train the SOMs. A cluster of genes (cluster of red signatures) in the lower half of the heat map identifies many of the fragile seed nodes. This cluster is the same set of fragile genes identified using dynamical computational analysis and RNA sequencing. Figure 5b shows the SOM distance matrix, a representation that allows for visualization of the high-dimensional spaces defined by the genes and their characteristics. The map is organized into clusters of similar color and topological space. Individual red lines connect neighboring nodes of the SOM and the colors in the regions containing the red lines indicate the distances between nodes with lighter colors representing shorter distances. **Supplementary Figure 18** shows the PCA plot as projections of the data set to the subspace spanned by the two eigenvectors with greatest eigenvalues. Individual component planes and U-matrices identify clusters of genes and the PCA projections associate the features that contribute most to the gene clusters. Figure 5c shows the number of genes associated with each cluster with the fragile gene cluster outlined. Fragile genes were most dissimilar to the other genes (robust genes) and form their own cluster. The SOMs, therefore, efficiently separate clusters of fragile and robust genes based on their genomic characteristics (SNPs, CNVs, CpG counts, etc.). Fragile gene clusters are most closely associated with increased frequency of SNPs, indels, simple repeats, DNAse hypersensitivity sites and novel sequence insertions. We then evaluated the biological function of the most fragile genes that clustered together in the SOMs. The most fragile genes from SOM cluster analysis were multiple genes involved with calcium handling and excitation-contraction coupling (RYR2, CACNA1C, TRDN, CACNA2D1), the rapid delayed rectifier K+ current (KCNQ1, KCNH2) and Na^+^-Ca^2+^ exchange current (ANK2 and SLC8A1).

**Figure 5.**
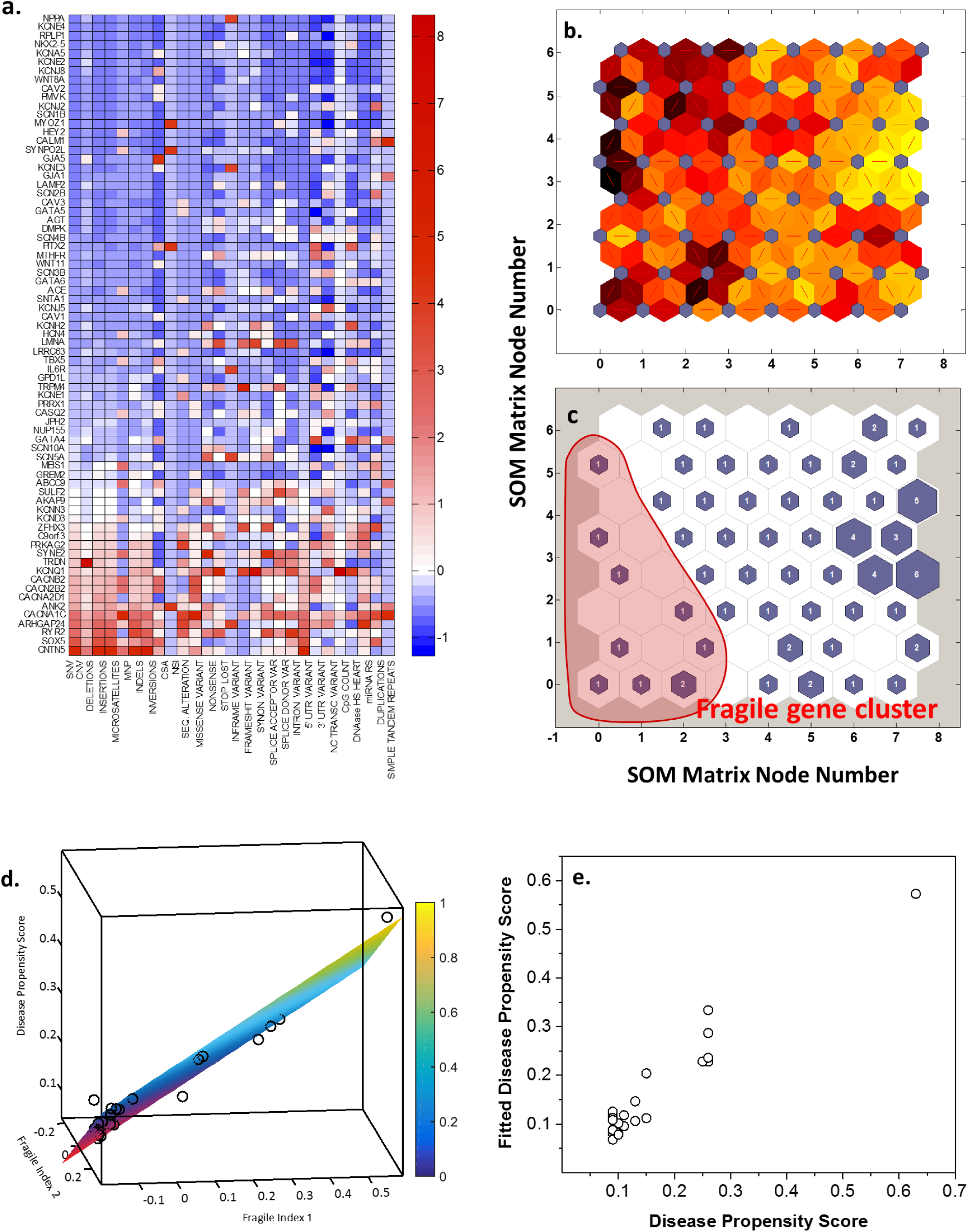
Variants in genes associated with fragile responses leads to disease. Unsupervised learning was used to understand the complex genetic architecture of arrhythmias including the sources of individual genetic variation that explain phenotypic variation. A density matrix was constructed using 28 different genomic features to describe known arrhythmia genes (seed nodes from AF, QTopathies, Brugada, CPVT and PCCD networks) and separately for each individual network gene. Genes were then clustered based on similar genomic features using the self organizing map (SOM) approach. **(a)** Heat map of normalized genomic feature values for all seed nodes based on the density matrix used to train the SOM. **(b)** Self-organizing map unified distance matrix (U-matrix) constructed using genomic features of seed genes. Individual red lines connect neighboring nodes (neural network) and the colors in the regions containing the red lines indicate the distances between nodes (lighter colors shorter distances). The U-matrix provides a simple way to visualize cluster boundaries on the map. **Supplementary figure 17** shows the individual weight matrix SOMs for each of the features. **(c)** Self organizing topology map with the number of training data vectors associated with each cluster. The fragile gene cluster is outlined in red. See **supplementary figure 18** for principal component plots of the genes and features. **(d)** Three dimensional scatter plot of fragile seed gene indices and their propensity for disease with best fit linear surface (R^2^ = 0.96). Partial least squares regression was used to construct a model using genomic features as an index of fragility and dynamical failure scores from action potential simulations as propensity for disease. Three PLS components were used to explain > 90% of the variation in genetic architecture. **(e)** Scatter plot of the observed disease propensity score and fitted disease propensity score from the 3 component PLS model (R^2^ = 0.94 for best linear fit).

Partial least squares (PLS) regression analysis was used to construct a model that relates genomic architecture and fragility to disease propensity. Disease propensity is the likelihood of developing an arrhythmia based on the failure of propagating an action potential. The PLS model is an independent model that incorporates quantitative information obtained from our dynamical analysis and genomic characteristics to predict disease propensity. Figure 5d shows the relationship between the fragile indices of seed node genes and their disease propensity with the best fit linear surface response shown as a plane (R^2^ = 0.96). The x and y axes, labeled as fragile index 1 and 2, are the two scores from the PLS regression analysis which together capture >85% of the variation contained in the 28 different genomic features. The z axis, labeled as disease propensity, represents the failure scores for individual genes derived from the dynamical ODE simulations (Figure 3g-h). Figure 5e shows the actual disease propensity score vs the fitted disease propensity score from our model (R^2^ = 0.94) and confirms the three component PLSR model predicts the response well.

The SOMs identified and confirm novel fragile genes and the genetic architecture associated with fragility. By looking at the biological function of fragile genes we gain insight to potential disease mechanisms. For example, specific genes involved in metabolism, cellular signaling (e.g., Wnt, Notch, Ca^2+^), histone and chromatin modification and regulation of RNA splicing also cluster with fragile channel genes. Several of our novel pathway findings were also supported by the literature. For example, multiple clinical trials suggest that histone deacetylase inhibitors and chromatin modulation influences cardiac electrical activity as seen by increased frequency of QT interval prolongation, T wave flattening and atrial fibrillation (38). HDAC inhibition also has been recently shown to reduce atrial fibrillation and remodeling in dogs (39).

We were particularly interested in a specific set of novel fragile genes involved in propanoate and butanoate metabolism that were predicted to influence arrhythmogenesis. At least one study associated inborn errors of metabolism caused by deficiency of propionyl CoA carboxylase with electrophysiological changes including prolongation of the QT interval (40) and two cases report children with propionic acidemia leading to arrhythmias with fatal outcomes (41, 42). Propanoate metabolism genes possess unique biological functions, network metrics (avg. shortest path length) and also associate with the fragile genes using unsupervised learning methods. Perhaps most importantly, the expression levels of propanoate metabolism genes were significantly different in human subjects with arrhythmias compared to subjects without arrhythmias (Figure 3a). Their *in-vivo* change in variance in human ventricular tissue also suggests that perturbation of their expression level may contribute to disease (Figure 4k). These metabolic genes are crucial in amino acid, fatty acid and cholesterol metabolism and many are expressed in the heart and muscle tissues (acetyl-CoA carboxylase), mitochondria (propionyl-CoA carboxylase) and brain tissue, thus contributing to our interest in evaluating their possible link to cardiac disease (43). In addition to the human RNA sequencing data that supports involvement of propanoate metabolism in arrhythmias, we sought to further experimentally test the importance for cardiovascular function using a series of zebrafish knockdown experiments. Morpholinos were used to knock down the propanoate metabolism genes SUCLG1/2, ACACA, PCCA and EHHADH and cardiac phenotypes in zebrafish were characterized using automated microscopic techniques. **Supplementary Figure 19** shows a combined box and whiskers and scatter plots of zebrafish heart rates and cardiac output for wildtype and propanoate metabolism gene knockdowns. Decreased contractility and morphologic changes in zebrafish hearts compared to wild type were observed when individual propanoate metabolism genes were knocked down. An apparent decrease in contractility was appreciated when SUCLG1/2 and ACACA genes were knocked down while a more subtle yet noteworthy change was also observed with EHHADH. Furthermore, quantitative decreases in heart rate and cardiac output were measured when all gene products were altered. Representative images of wild type and ACACA knockdown zebrafish hearts are shown in **supplementary figure 19**. Qualitatively, these images show knockdowns with morphologically distinct hearts and increased pericardial edema indicative of impaired cardiac activity.

## Discussion

It is now widely accepted that most proteins function as components of networks, and considering their activity in the context of the networks they belong to is useful for understanding how individual protein activities contribute to network based emergent physiological functions. As a network node that is required for dynamic function, a protein within a network needs to be defined by at least two characteristics, its quantity and its ability to interact with its partners. These two characteristics are also the key parameters for the dynamical models: initial concentration of reactant and reaction rate. Thus it should be relatively simple to convert a directed sign-specified network into a system of coupled differential equations and thereby develop a systems level understanding of why some genetic constructs but not others are correlated with disease phenotype. However, our knowledge of protein concentrations and reaction rates is currently sparse, and it is not possible to build reliable dynamical models directly from large network representations. New data integration methods are needed to relate statistical and network representations to parameter values in dynamical models, and this study represents a first step in that direction.

When starting the study we had thought that disease determinants would be associated with changes in robust kinetics outside the boundaries of homeostatic tolerance and that these determinants would primarily be coding variants. However, we found that specific types of both non-coding and coding variants in arrhythmia genes were associated with fragile kinetics and correlate with disease propensity. Importantly, our studies show that these multigene effects arise because of the association with fragile kinetic parameters of cardiac electrophysiology. These results were unexpected, based on what is known for common cancer mechanisms such as mutations in receptor tyrosine kinases and GTPases. We would have expected that robust kinetic parameters of gene products could have been associated with overrepresented regions of structural variation in the genome associated with arrhythmias. In fact, others have even suggested that because of evolutionary selection constraints, it is just as likely that most genes involved in common polymorphic rearrangements are tolerant of changes in certain types of architectural variation (44). The power obtained by integration of multiple mathematical approaches and human RNA sequencing has enabled us to obtain a counterintuitive and deep understanding of genotype-phenotype relationships and improved our ability to predict disease risk.

The data from multiple modeling studies, human RNA sequencing as well as the experiments in the zebrafish model together support the hypothesis that the genetic architecture of fragile electrophysiological responses and arrhythmias is responsible for increased disease propensity. The self-organizing maps efficiently classify many genes as fragile and demonstrate substantial enrichment for genomic regions associated with cardiac electrical parameters over the remainder of the genome. This is a new approach that provides a comprehensive framework for understanding how variations of a convergent phenotype may manifest from multigene interactions. Similar to other multigenic disorders that were initially characterized as being monogenic, pathophysiologies of cardiac electrical activity are likely to be caused, modulated or suppressed by allelic heterogeneity and mutations at multiple loci that are associated with fragile kinetics (45). Further mechanistic understanding of how the multiple genomic loci interact will always come from studies focused on physical and functional interactions or regulation of the proteins involved. This study provides direct evidence in support of this assertion. It should be noted that the partial least square based correlation approach between kinetic parameters and genomic loci has its limitation in providing comprehensive mechanistic understanding. Nevertheless defining such relationships provides the boundary conditions within which the mechanisms will operate.

The network based approach for prediction of additional genomic loci and testing of the predictions in humans and the zebrafish models indicates that there are likely to be additional genomic loci that are related to the origins of arrhythmias and disease progression or that contribute to disease propensity in the context of other genomic variations or environmental effects. A collaborative study of Noonan syndrome allowed one of us to use a pathway based approach to identify a gene that turned out to be disease related upon subsequent laboratory based analyses of patient samples (46). Perhaps further studies on the genomes of patients with arrhythmias could lead to the identification of additional genes involved in the pathophysiology of cardiac electrophysiology and provide a basis for developing new drugs to treatment of arrhythmias. We suspect that the novel relationships identified in this study between fragile kinetic parameters and architectural variation could be a common theme for many diseases with common complex traits. For example, autism, schizophrenia, epilepsy, Parkinson disease, Alzheimer disease, immunological disorders and emphysema, among others, have all been shown to result from structural variation in the genomes of at least some patients (44). The challenge will be to define kinetic parameters and identify models capable of recapitulating the relationship between fragility in dynamics and genomic determinants as one cause in multiple complex diseases.

## Acknowledgements

This research was supported by NIH grants P50-GM071558, R01-GM54508 and R01-HL109264. TK was also supported by a fellowship from the Sarnoff Cardiovascular Research Foundation.

